# Sandalwood, an Indian medicinal plant attenuates the microbial growth and influence up/down regulation of the metabolites

**DOI:** 10.1101/2021.03.01.433331

**Authors:** Romana Parveen, Tooba Naz Shamsi, Sadaf Fatima

## Abstract

The methanolic extract of sandalwood (SwME) was prepared by soxhlet apparatus and the antibacterial assay was performed. Further, the metabolite profiling of SwME and lysates of *E. coli* and *E. coli* grown in the presence of SwME was generated. SwME showed maximum inhibition against *E. coli* (MTCC 443) i.e. 82.71%, and minimal against *B. subtilis* (MTCC 736) i.e. 26.82%. The metabolome profiles of *E. coli* and SwME were generated using gas chromatography-mass spectrometry (GC-MS) technique. Comparative studies were done to understand to what extent metabolite modifications differ between SwME, *E. coli* lysate and the *E. coli* strain grown in presence of extract. Result revealed 23 peaks with major compounds present in *E. coli* were 9-Octadecenoic Acid (Z)-, Methyl Ester (26.85%), Hexadecanoic Acid, methyl ester (20.5%) and Hexadecanoic acid, trimethylsilyl ester (15.79%). When *E. coli* was grown in the presence of SwME, GC-MS analysis showed 25 peaks with major compounds such as 9-Octadecenoic Acid, Methyl Ester (21.97%), Hexadecanoic Acid, Methyl Ester (17.03%), and Hexadecanoic Acid, Trimethylsilyl Ester (14.96%). Correlating the metabolic profiles with the changes occurring is essential to progression their comprehension and in the development of new approaches to identify the metabolomics regulation in *E. coli* in response to SwME.

## 1. Introduction

With the advancement in human livings effected by technology and such innovations, they say our disease causing organism have also developed themselves. This is currently the most focused aspect of humans, the new or improved ways of defense and cure from the life threatening disease. The present chemical based pharmaceutical is looking new hopes in traditional or natural origin medicines which would be efficiently effective against disease causing microorganism which are now said to immune themselves from the chemical based medicine available in the market (Baker et. al. 2007, Beghyn et. al. 2008 and Brinskin 2000). Nature has all the solutions to mankind problems; such is the natural or ayurvedic medicine derived from the plants, animals or marine. Therefore, several plants have been selected based on their use in traditional systems of medicine (Ji et. al. 2009). Among the thousands of such therapeutic plants, Sandalwood is one of the well-known plants from the Indian origin that has been used in cosmetics and therapeutics and is used by Ayurvedic medicinal practitioners in India for almost 3000 years. It is known for its multi-therapeutic properties and is being used since a long time in this part of world as a cosmetics and therapeutic agent. The Indian plant, *Santalum album* may serve as a dietary antioxidant, with several modes of action to protect the body organs against oxidative damage and age-related cognitive decline. Aqueous extract of *Santalum album* showed good activity against *Staphylococcus aureus, Bacillus subtilis, Pseudomonas aeruginosa* and *Escherichia coli* and the aqueous extract showed maximum activity against *S. aureus* (Shamsi et. al. 2014).

Plants are known to possess the various complex phytochemical compounds that may be used in the prevention and treatment of several human diseases (Hall et. al. 2002). Identification of phytochemicals or bioactive compounds and their characterization are required to explore the medicinal value of the plant which is an ongoing challenge in this era. This process needs the several extraction methods for the metabolites extraction and the advanced techniques that can identify, detect and characterize these bioactive compounds (Wolfender et. al. 2015). Metabolomics is a field where various analytical strategies and platform in application of natural extracts develop the chemical profiles. The information obtained from this approach might be useful in the determination of various biological aspects of plants such as defense, growth, development, productivity, responses to external stresses etc. (Desai and Alexander 2013, Misra et. al. 2014). Gas chromatography (GC) coupled with mass spectrometry (MS) is the analytical platform and strategy that can be employed for the analysis of metabolites found in the plant extracts qualitatively and quantitatively (Wong et. al. 2015). GC-MS analysis can identify up to approximately 100 metabolites in the simple plant extracts and can generate a metabolite profile (Benina et. al. 2013, Obata et. al. 2013, Sauter et. al. 1991, Lee et. al. 2014). The effective and better identification of the obtained metabolites can be easily done by comparing the results with the database stored in the large mass spectral library based on GC-MS electron impact (EI) (Musharraf et. al. 2013). The sample preparation for the extraction of metabolites also plays a key role in this analytical platform of metabolomics and is known to be a major factor that affects the profiling of metabolite (Ying et. al. 2009). Metabolites extraction can be done in various organic solvents such as ethanol, water, methanol, and hexane and can be carried out by using various conventional extraction methods followed by lyophilization and then monitoring and generating the profiles of extracted metabolite using several analytical platforms (Lapornik et. al. 2009, Ahmad and Shah 2015). Various factors such as type of solvent, extraction method, temperature etc. affect the metabolic content of crude extracts of plants (Kindt et. al. 2009). For the study of above mentioned properties of *Santalum album,* antibacterial assay and GC-MS analysis was performed to know the metabolite composition of *Santalum album* methanolic extract, *E. coli* (the most susceptible bacterial strain) lysate and the *E. coli* grown in presence of *Santalum album* methanolic extract.

## 2. Materials and methodology

### 2.1. Plant material/Chemicals

The medicinal plant sandalwood (*Santalum album*) was purchased from Local Ayurvedic Clinic, Rampur, Uttar Pradesh, India. The chemicals of analytical grade were purchased from Merck and HI media.

### 2.2. Test microorganism

The identified bacterial strains *Staphylococcus aureus* (MTCC902), *Escherichia coli* (MTCC443), *Bacillus subtilis* (MTCC736), and *Pseudomonas aeruginosa* (MTCC2453) were obtained from NCCS, Pune, India. Stock cultures of all the tested bacterial strains were maintained on nutrient agar slants at 4°C.

### 2.3. Preparation of methanolic crude extract of sandalwood

The sandalwood was ground finely in a grinder mixer and then 10 g of powdered sandalwood was placed in a soxhlet apparatus. Extraction was carried out with 250 ml of methanol for 36-48 h. The temperature was maintained below the boiling point of the solvent. Further, the extract was filtered through a 0.45 μm filter. The filtrate was then concentrated by Rotary Evaporator and it served as the methanol extract of sandalwood (SwME) for the further exploration and activities. The extract was stored in a refrigerator at 4°C for further use.

### 2.4. Determination of antibacterial potential of sandalwood methanolic extract (SwME)

SwME was evaluated for its antibacterial potential against several bacterial strains as the method described by Shamsi et al. 2014 with some modifications. A single colony form agar plates of each bacterial strains was inoculated in autoclaved Luria broth (LB) media for bacterial growth. The cultures were incubated at 37°C for 12-24 h. Next day, the LB broth was added to the bacterial cultures to dilute it up to 1×10^4^ colony-forming units (CFU). Approximately 50 μl of 5mg/ml SwME (250μg SwME) was mixed with diluted bacterial cultures in 96-well plate. The final volume in each well was maintained up to 300 μl using LB media. Ampicillin (Stock 10 mg/ml) was used as positive control and 25 μl of it (250 μg ampicillin) was applied in each well containing LB media and bacterial broth culture. The results were compared with negative controls which contained the LB media and bacterial culture. Blanks contained LB media only and the LB media with SwME. 96-well plate was further incubated at 37°C for 14-16 hrs. After incubation, absorption measurements were determined using ELISA microplate reader at 600 nm. The percentage mean growth inhibition (MGI %) was calculated by % MGI = [(A_c_ – A_t_)/A_c_] x 100.

Where, A_c_ and A_t_ are the absorbance of negative control and SwME treated strains respectively.

The EC_50_ value (concentration causing 50 % reduction in the bacterial growth) was calculated using Microsoft office Excel 2007, based on the readings.

### 2.5. The GC-MS analysis

#### 2.5.1. Homogenization and extraction

Homogenization and extraction were the preliminary step to quench biomass for the release of metabolites. Homogenization is carried out by a grinder. The solvent methanol was degassed and cooled before the extraction procedure.

#### 2.5.2. Biomass Quenching

SwME, and the lysates of *E. coli* and *E. coli* grown in the presence of SwME were taken and the biomass was centrifuged for 20-30 min to pellet down the biomass at 5000 rpm at 4°C. Then, the supernatant was discarded and an approximately 1 ml aliquot was preserved for the determination of leakage of internal metabolites. Further, the pellets and supernatants were frozen in liquid nitrogen followed by storing them at −80°C and then the further analysis was done.

#### 2.5.3. Extraction of metabolites

The biomass in the form of pellets were mixed with 500 μl of 100% chilled methanol (−48°C), followed by freezing in liquid nitrogen and then allowed to thaw on dry ice. The process of freezing followed by thawing was performed several times which helped in the permeability of the cell membrane, which resulted in the metabolite leakage from the cells. The samples were then centrifuged at 10000 g for 5-10 min at 4°C. Then, the supernatant was obtained and preserved on dry ice. Approx. 500 μl aliquot of 100% methanol (−48°C) was further mixed with the pellet and then stored on dry ice to perform further analysis.

#### 2.5.4. Mass spectrometric analysis

GC-MS analysis of lysate of *E. coli,* SwME and *E. coli* grown in presence of SwME was performed on a GC-MS equipment. The dimensions of GC-MS column (non-polar column) used were about 30 m ×0.25 mm with a film thickness of 0.25 mm based on capillary standard. The flow rate was maintained up to 0.5 ml/min and approximately 1μl of the concentrated sample was utilized. The methanolic samples were run fully at a range of 50-650 m/z and the inlet temperature was maintained up to 250°C. The samples were run for approx. 90 min and GC-MS was examined by using electron impact ionization at 70eV. The obtained spectrums of the samples were compared with the database of spectrum of known compounds which are stored in the GC-MS NIST library to evaluate the total ion count for compound identification and robust quantification. Further, the peak areas, retention time etc. were measured and data processing was performed by Turbo-Mass software. The raw GC-MS chromatograms were transferred to a server station by recording two scans per second in full scan mode.

#### 2.5.5. Data acquisition by GC-MS

Peak identifications were carried out by matching retention indices and mass spectral similarity against a user-defined metabolite library. Metabolite peaks were quantified by area of target ion traces. The obtained number of metabolites was counted per chromatogram.

#### 2.5.6. Identification of components

Identification of the obtained compounds were done by comparing their GC-MS mass spectrum by using the database of National Institute Standard and Technology (NIST) with software Turbomas 5.2 which stores approximately 62,000 patterns which was based on their molecular mass, molecular structure, calculated fragments. The database then generated a table which included the name, molecular weight and structure of the tested components by comparing to the database. The % relative of each compound was calculated by comparing the average peak area in the chromatogram to the total areas. The detection of compounds helped in the determination of the current commercial, industrial and traditional use of crude extracts as an herbal medicine. This method also helped in the determination of the best extraction methods for these components. The identified compound now further can be explored for their putative biological or therapeutic relevance. GC-MS software that was able to perform multi-target analysis also included quantitation by area and height of the peak in the chromatogram.

### 2.6. Statistical analysis

The experiments were performed in triplicates (n=3). Graph Pad Prism Software was used to prepare Graphs and to calculate the statistical parameters such as mean, standard error of mean (SEM) etc. The data obtained were expressed as ‘mean±SEM’.

## 3. Results

### 3.1. Determination of antibacterial activity of SwME

Antibacterial potential of the SwME was evaluated by assessing the percentage mean growth inhibition in the presence of SwME and the results were compared to the proper positive control. The results exhibited fair antibacterial property *in-vitro* against all the tested strains i.e. the bacterial growth was diminished in the presence of SwME as shown in Fig. 1. Results showed that SwME had strongest inhibitory activity against *E. coli* (MTCC 443) i.e. 82.71% and minimal inhibition against *B. subtilis* (MTCC 736) i.e. 26.82% at approx. 0.833 mg/ml concentration of SwME. Results were quite comparable with synthetic antibacterial drug ampicillin which showed almost complete inhibition of bacterial pathogen at the same concentration of antibiotic.

**Figure 1:**
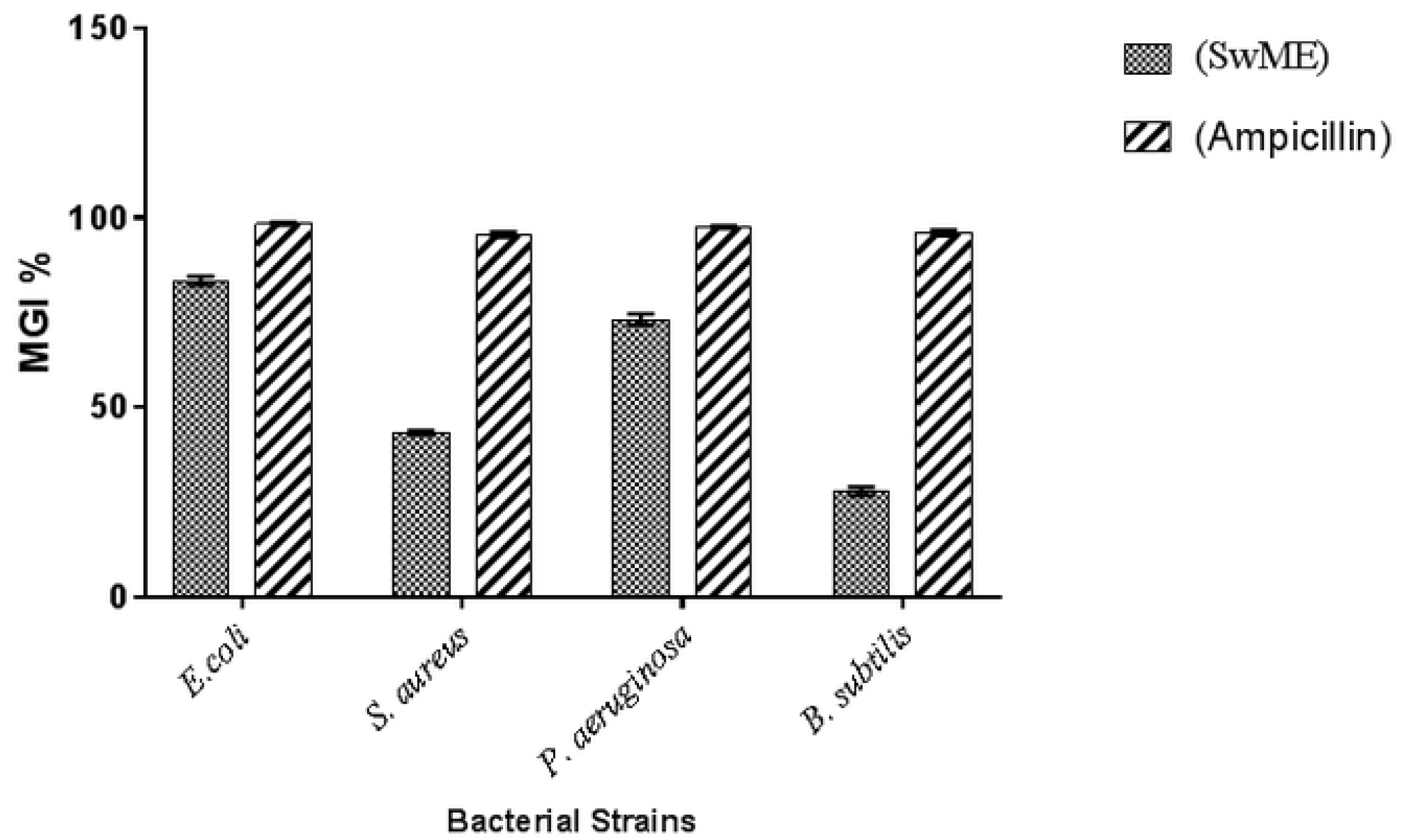
Bar diagrammatic representations of *in vitro* antibacterial activity of sandalwood methanolic extract. The bars represent the percentage mean growth inhibition by methanolic extract of sandalwood when tested against 4 bacterial strains.

Generally, *B. subtilis* was the only strain which was less susceptible to the antibacterial effect of the SwME and ampicillin as compared to the other strains. SwME exhibited different degree of growth percentage inhibition among tested bacterial strains which were presented in the form of EC_50_ values (Table 1).

**Table 1:** Determination of EC_50_ values of SwME and ampicillin and represented in mg/ml.

| Bacterial strains | EC<sub>50</sub> (mg/ml) |
| --- | --- |
| *E. coli* (MTCC 443) | 0.50 |
| *S. aureus* (MTCC 902) | 0.95 |
| *P. aeruginosa* (MTCC 245) | 0.57 |
| *B. subtilis* (MTCC 736) | 1.54 |

### 3.2. GC-MS analysis

Identification and quantification up to 100 metabolites can be performed using an important and critical technique i.e. GC-MS based metabolite profiling. These metabolites may incorporate sugars, hydrocarbons, phenolic compounds, carboxylic acids, sugar alcohols, alcohol acids, esters, amino acids, and polyamines etc. and provide a far reaching scope of the primary metabolism central pathways. The present study emphasizes on use of the GC-MS analysis for identification and quantification of the metabolites or compounds present in the methanolic extract of sandalwood, *E. coli* and the *E. coli* grown in presence of SwME. The GC-MS analysis spectra of SwME, *E. coli* and *E. coli* +SwME showed and identified the presence of several phytocompounds that could contribute to the ethno-medicinal quality and the pharmacological nature of the plant. Phytocompounds were identified and confirmed by comparing their retention time, peak area, molecular formula etc. to the database of NIST library.

#### 3.2.1. Identification of compounds in SwME

GC-MS chromatogram of the SwME (Fig. 2) showed 43 peaks indicating the presence of forty one compounds. The chemical compounds identified in the methanolic extract of the sandalwood extract are presented in Table 2. Based on abundance, the most prevailing compounds present in the methanolic extract were Piperidine, 1-[5-(1,3-Benzodioxol-5-yl)-1-oxo-2,4-Pentadienyl] (15.08%), 2-Penten-1-ol, 5-(2,3 Dimethyltricyclo[2.2.1.0(2,6)]Hept-3-yl)-2Methyl (14.44%) and 2-Penten-1-ol, 2-Methyl-5-(2-Methyl-3-Methylene-2-Norbornyl) (8.83%). Acetonyl Decyl Ether (0.09%), Pregnane, Silane Derivative (0.1%) and 1, 2-Benzenedicarboxylic Acid, Diisooctyl Ester (0.2%) were present in low amount. The result of GC-MS analyses unveiled that the SwME is primarily composed of oxygenated hydrocarbons and prevalently phenolic hydrocarbons. The phytochemical compounds revealed through GC-MS analysis are those phytochemicals which are responsible for various pharmacological activities such as antimicrobial, anti-inflammatory, antioxidant activities etc.

**Table 2:**
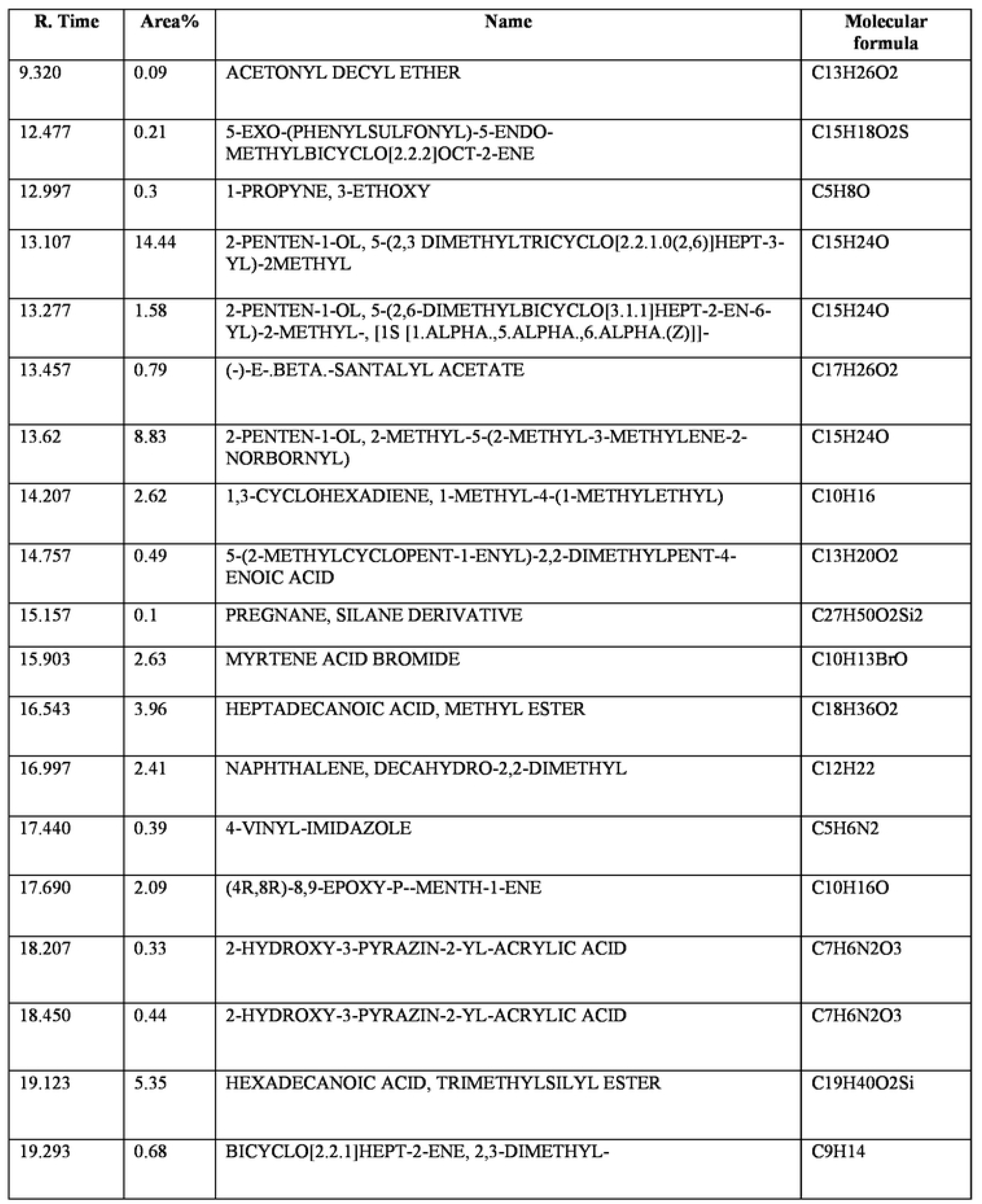
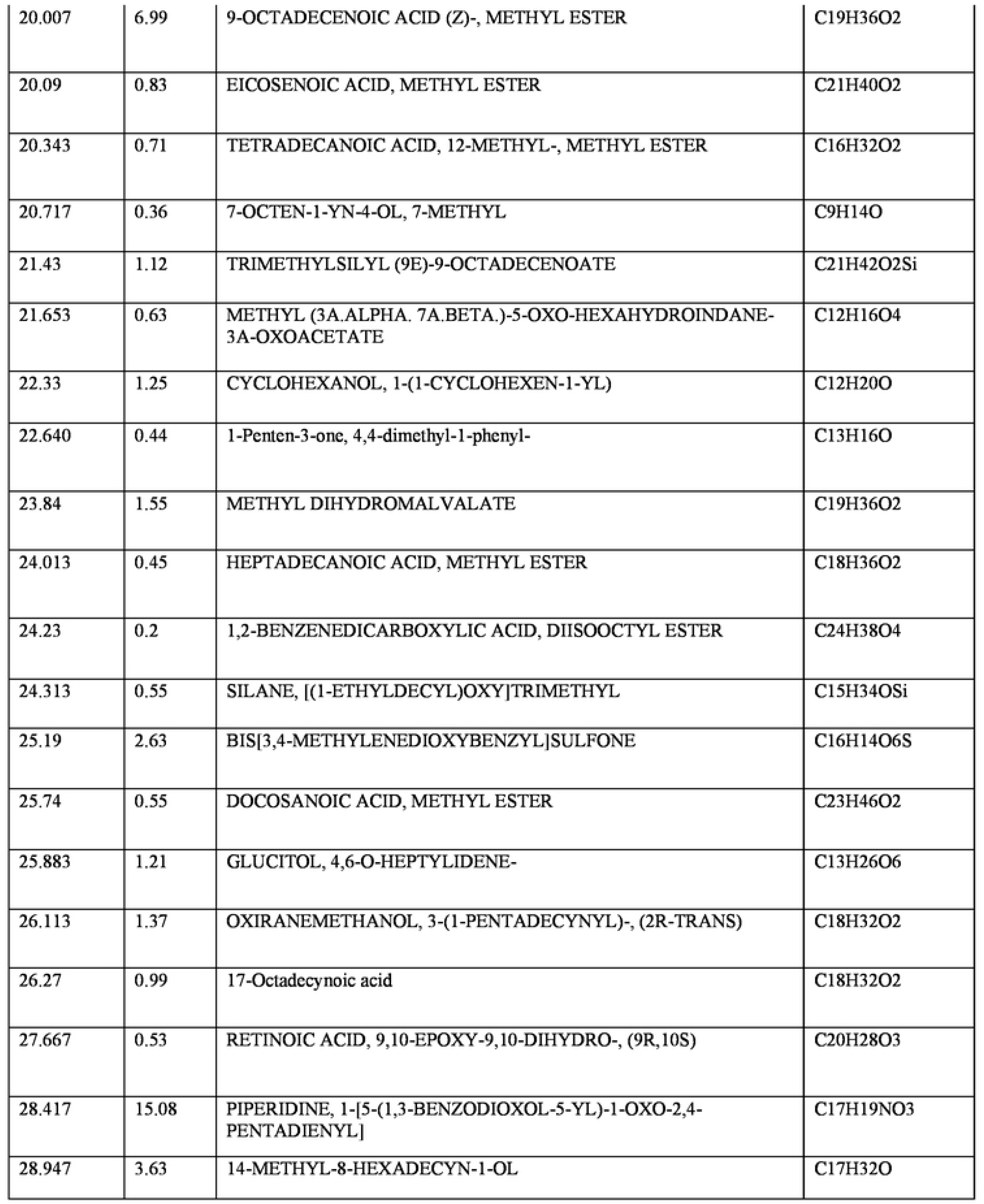
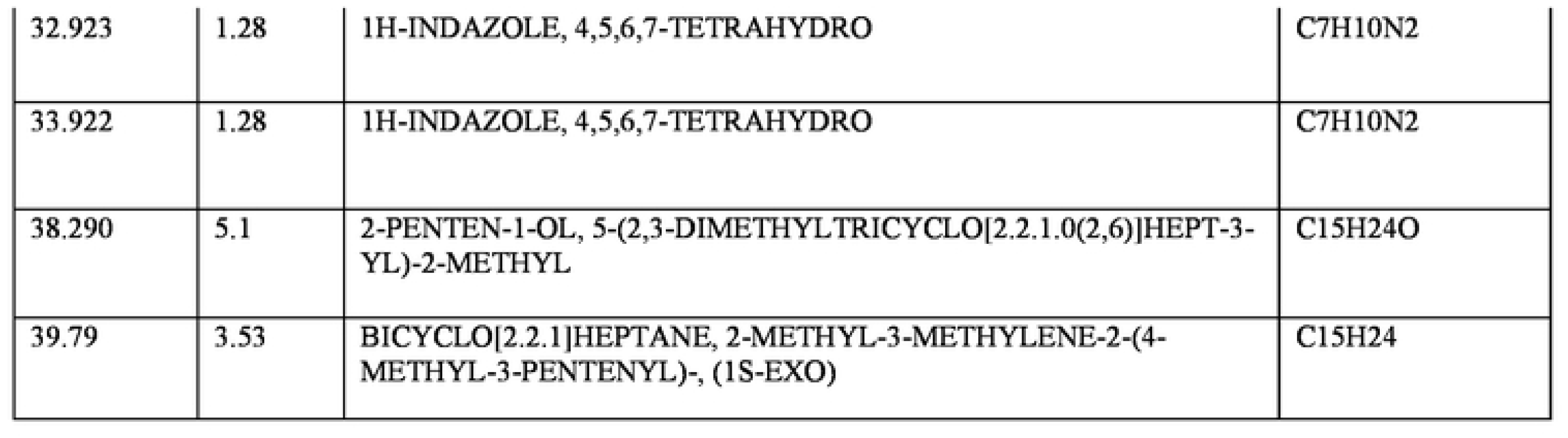
Biologically active chemical compounds of methanol extract from sandalwood, showing retention time, molecular formula and peak area (%).

**Figure 2:**
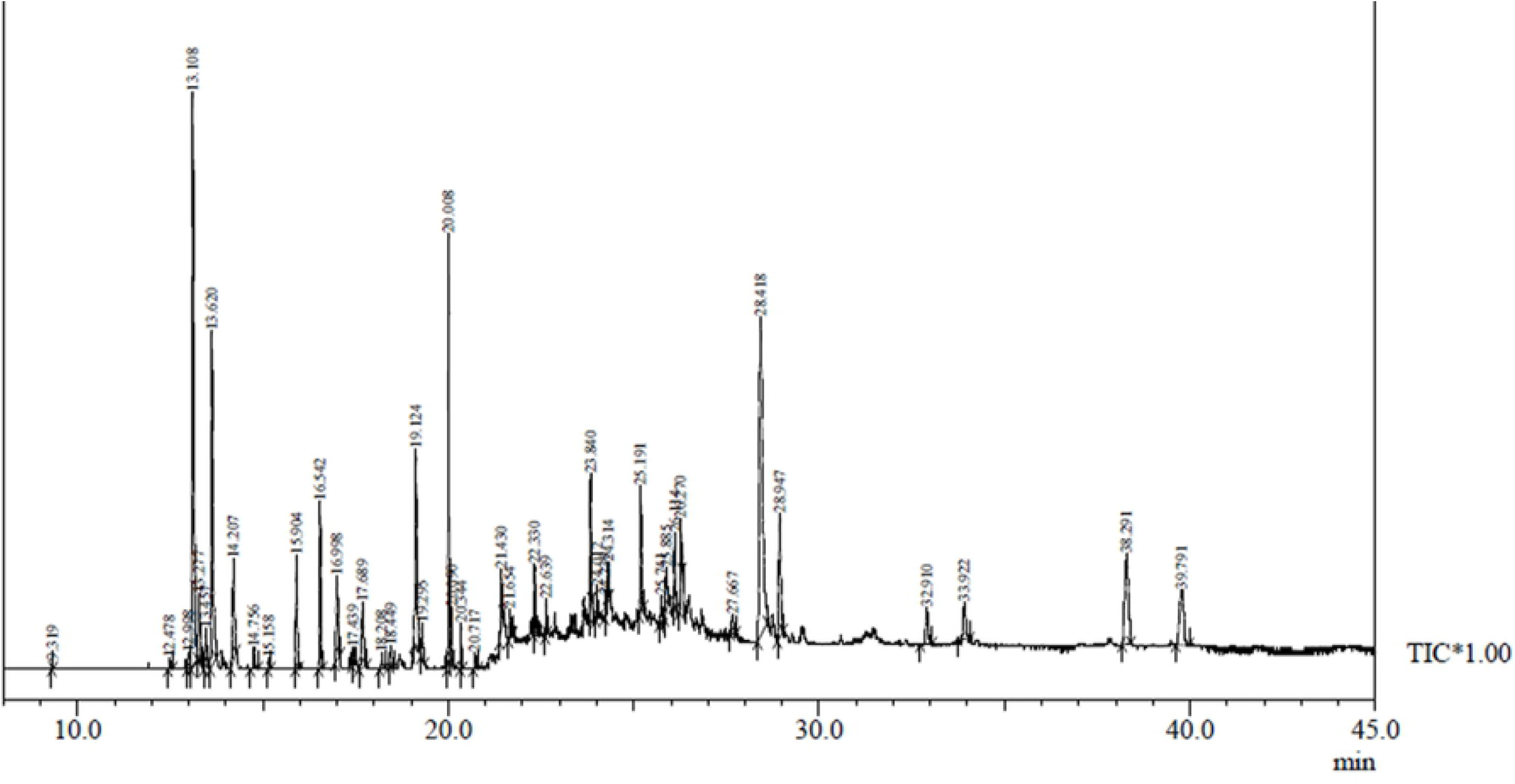
A typical chromatogram of the bioactive compounds present in methanol crude extract of sandalwood plant showing different peaks.

#### 3.2.2. Identification of compounds in E. coli lysate

Further, the lysate of *E. coli* revealed the presence of 23 peaks in the GC-MS chromatogram which indicates the presence of 20 compounds. The major compounds obtained were 9-Octadecenoic Acid (Z)-, Methyl Ester (26.85%), Hexadecanoic Acid, methyl ester (20.5%) and Hexadecanoic acid, trimethylsilyl ester (15.79%) and the 1-Hexyl-1-Nitrocyclohexane (0.23%), Pentadecane (0.32%) and Heptadecanoic Acid, Methyl Ester (0.35%) were found in rare amount (Fig. 3, Table 3).

**Figure 3:**
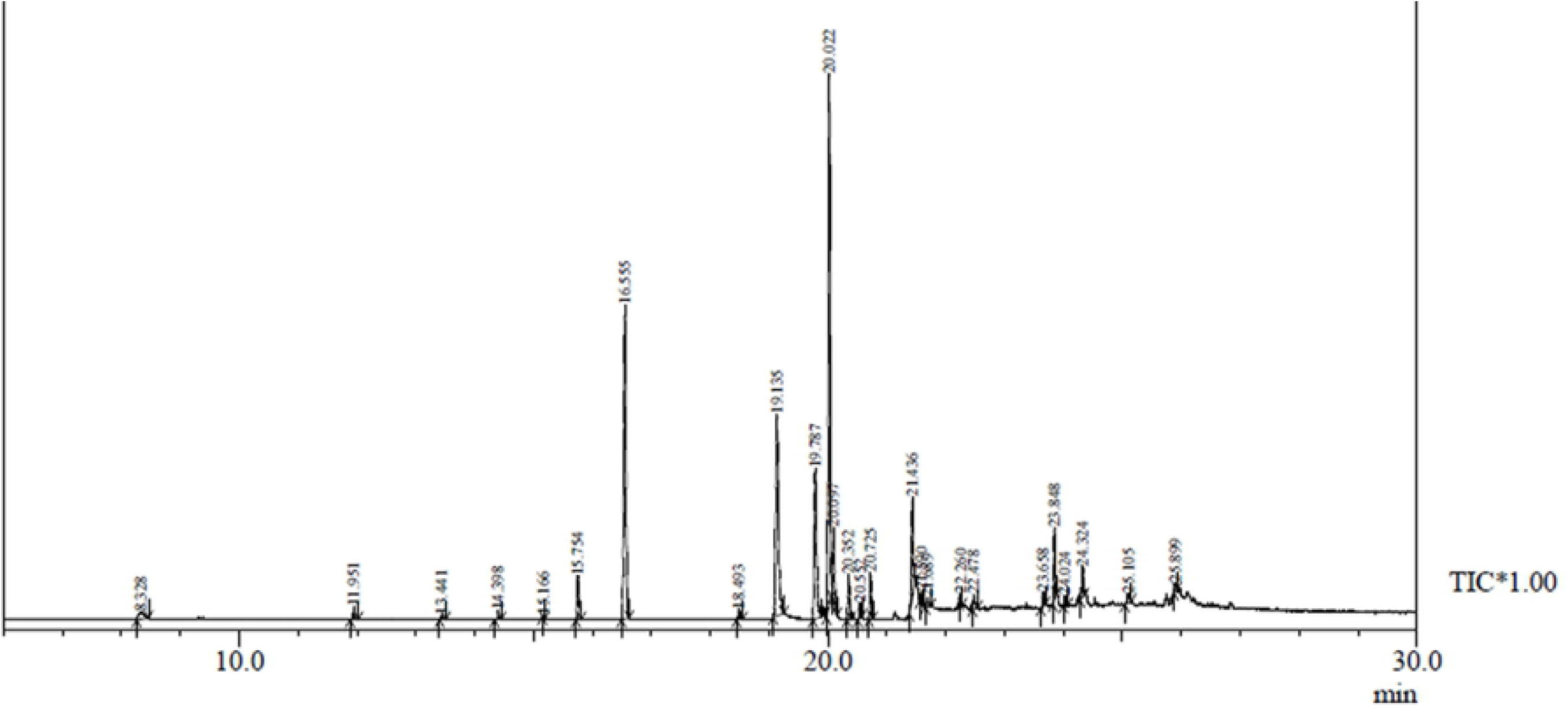
A typical chromatogram of the metabolites found in the methanolic lysate of *E.coli.*

**Table 3:**
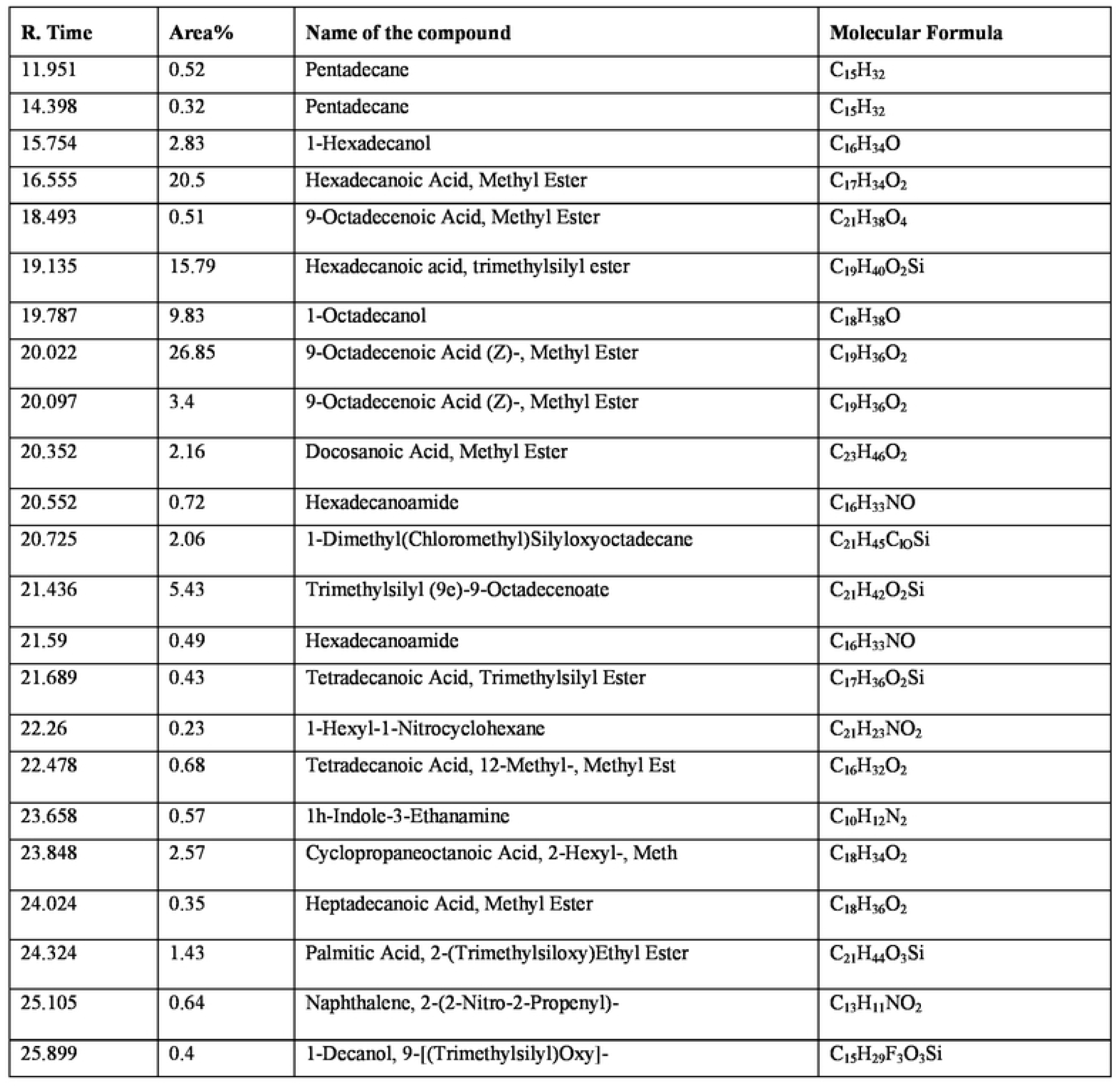
GC-MS analysis revealed the presence of compounds in the methanolic lysate of *E. coli* showing retention time, molecular formula and peak area (%).

#### 3.2.3. Identification of compounds in E. coli grown in the presence of SwME

While, when *E. coli* was grown in the presence of SwME to find the impact of bioactive compounds of the SWME extract on the metabolites present in the *E. coli,* the methanolic lysate was explored to find the up or down-regulated metabolites using GC-MS technique. The results obtained showed 25 peaks indicating the presence of twenty one compounds. This time, some different compounds were observed which was neither present in SwME nor in *E. coli.*

The results obtained here suggested that, sandalwood up regulated few metabolites in *E. coli* which may be present in the negligible amount and hence this up regulation might be responsible for the bacterial death. The different metabolites observed were 3-Ethyl-8-Methyl-2-Oxatetracyclo [4.4.0.0(1, 4).0(6, 8)] Decane, Benzene, Ethoxy, Trans-9-Octadecenoic Acid, Trimethylsilyl Ester, Oleic Acid, Trimethylsilyl Ester, Octadecanoic Acid, Trimethylsilyl Ester, 1-Hexyl-2-Nitrocyclohexane, Tetracosanoic Acid, Methyl Ester, 9-Octadecenoic Acid, 2-[(Trimethylsilyl)Oxy]-1-[[(Trimethylsilyl)Oxy]Methyl]Ethyl Ester, Silane, [(10-Chlorodecyl)Oxy]Trimethyl, 9,12-Octadecadienoic Acid (z,z)-, 2-[(Trimethylsilyl)oxy]-1-[[(Trimethylsilyl)Oxy]Methyl]Ethyl Ester etc. The major compounds were 9-Octadecenoic Acid, Methyl Ester (21.97%), Hexadecanoic Acid, Methyl Ester (17.03%), and Hexadecanoic Acid, Trimethylsilyl Ester (14.96%) and Tetradecanoic Acid, 12-Methyl-, Methyl Ester (0.35%), 1-Hexyl-2-Nitrocyclohexane (0.36%), 9,12-Octadecadienoic Acid (z,z)-, 2-[(Trimethylsilyl)Oxy]-1-[[(Trimethylsilyl)Oxy]Methyl]Mthyl Ester (0.38%) were found in minimal percentage. GC-MS analysis showed up/down regulation of various compound. Some compounds were turned out to be rare and some compounds get up regulated and showed high peak in the chromatogram (Fig. 4, Table 4).

**Figure 4:**
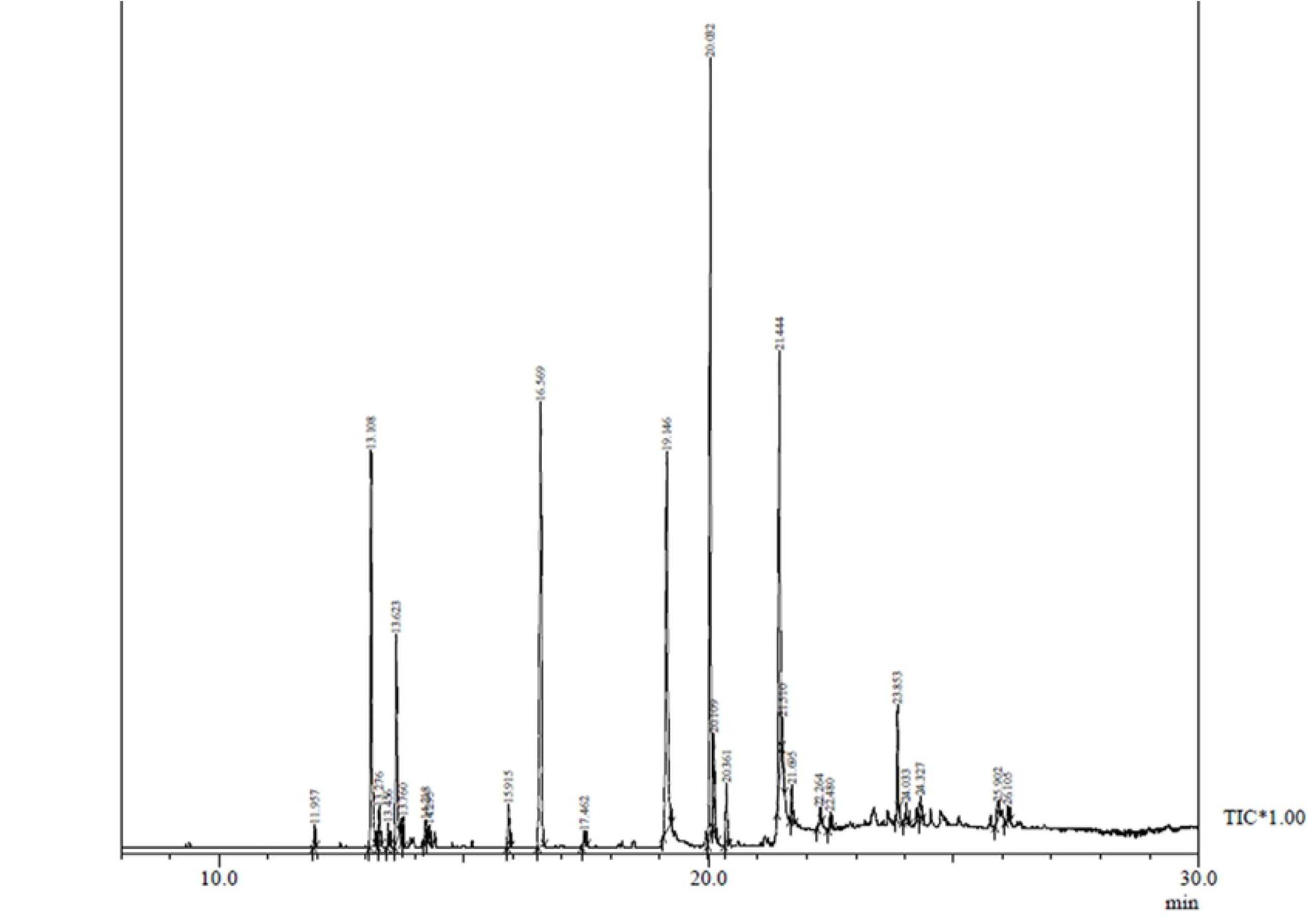
A typical chromatogram of the metabolites present the methanolic lysate of *E. coli* grown in presence of sandalwood methanolic extract.

**Table 4:**
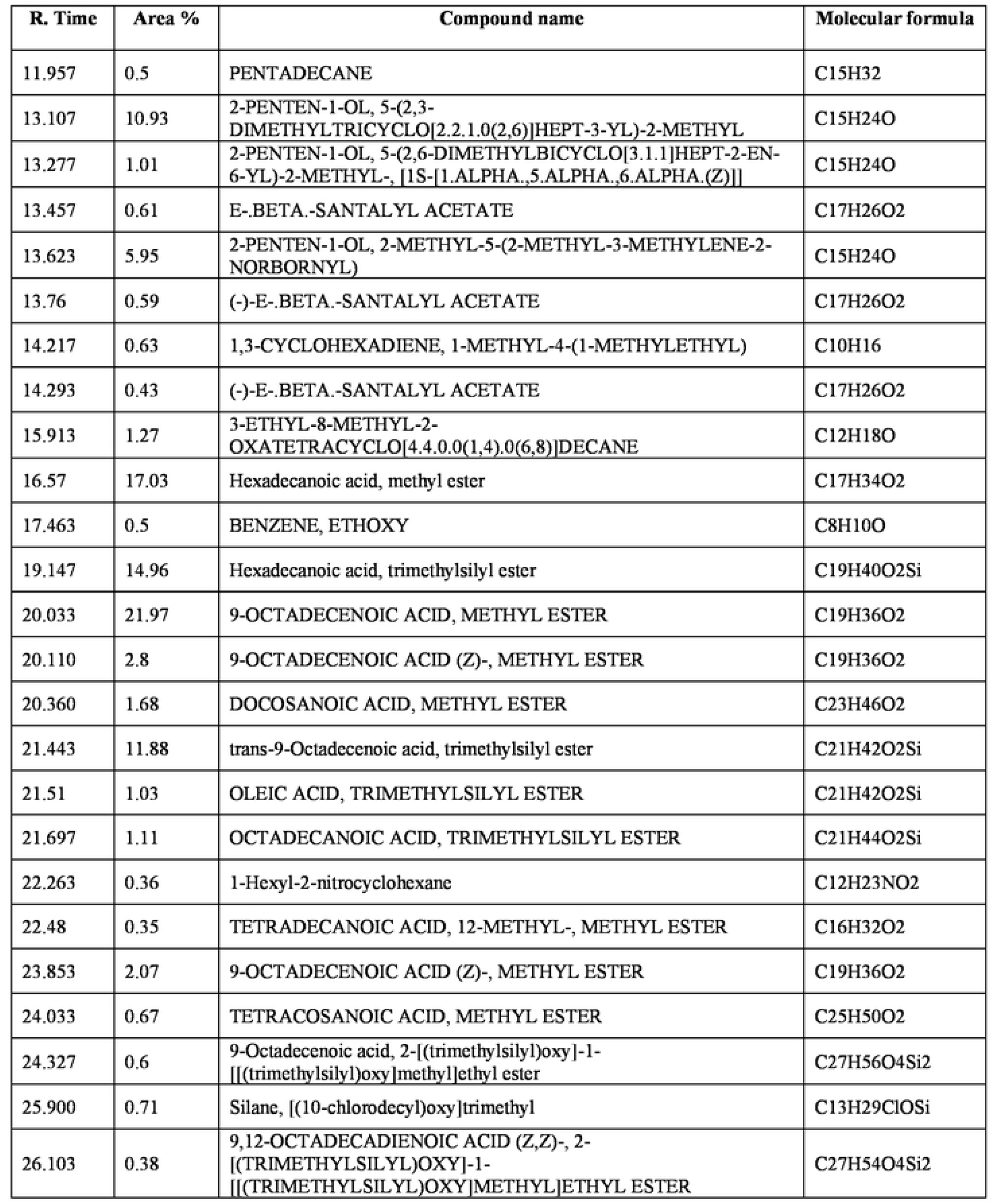
GC-MS analysis revealed the presence of compounds in the methanolic lysate of *E.coli* grown in presence of sandalwood methanolic extract showing retention time, molecular formula and peak area (%).

There are some phytocompounds such as Pentadecane (0.52%), 9-Octadecenoic Acid (Z)-, Methyl Ester (26.85%), Docosanoic Acid, Methyl Ester (2.16%), Tetradecanoic Acid, 12-Methyl-, Methyl Est (0.68%) found in *E.coli* lysate. These compounds get downregulated with peak area 0.5%, 21.97%, 1.68% and 0.35% respectively when grown in presence of SwME.

The present investigation is only a preliminary characterization of the presence of some particular properties in sandalwood extract. Further investigation and in-depth knowledge is required to identify the biochemical and phytochemical functions of the herbs. The metabolites which were identified here by GC-MS analysis revealed that they possess many biochemical and pharmacological potential and can be based on Dr. Duke’s Phytochemical and Ethnobotanical Databases created by Dr. Jim Duke of the Agricultural Research Service/USDA.

## 4. Discussion

Medicinal plants are considered new as new assets for producing agents that go about as alternatives to antibiotics against antibiotic-resistant bacteria. It is very important for an antimicrobial agent to be eco-friendly and biologically safe. This study revealed that sandalwood could be a promising antibacterial agent as it showed inhibition against various bacterial strains i.e. the maximum and minimum inhibition of SwME was found to be against *E. coli* and *B. subtilis* respectively. Also the compounds which get up/down regulated might be responsible for this activity. Shamsi et al. in their study investigated that the aqueous extract of *S. album* showed effective inhibitory activity against *S. aureus* (MTCC 902) (Shamsi et. al. 2014). The results from antibacterial activity of SwME revealed that *E. coli* was the most sensitive strain with highest % inhibition towards methanolic extract. (Stenz et. al. 2008) in their study showed when 9-octadecenoic acid was added after primary adhesion was developed, it stimulated biofilm formation and hence bacterial growth. Through our study, it is confirmed that when *E. coli* was grown in presence of SwME, 9-octadecenoic acid was down-regulated which confirms the result of the above study.

The presence of different compounds in SwME through GC-MS analysis showed sandalwood to be the promising therapeutic agents as the compounds present in methanolic extract possess antimicrobial, antioxidant, antiulcer, anti-asthmic activity etc. Hexadecanoic Acid present in methanolic extract of sandalwood possess Anti-inflammatory, Antioxidant, hypocholesterolemia nematicide, pesticide, anti-androgenic flavor, hemolytic, 5-Alpha reductase inhibitor, potent mosquito larvicide (Aparna et. al. 2012, Kumar et. al. 2010, Rahuman et. al. 2000, Miustapha et. al. 2016), 9-12 Octadecenoic Acid (Z)-, Methyl Ester shows anti-cancer and antimicrobial activity (Yu et. al. 2005, Rahman et. al. 2014). Piperidine has characteristic antiasthamic activity (Matsushita et. al. 1998). The literature survey also confirmed the presence of 9 Octadecenoic Acid as a plant metabolite in the hexane soluble extract of the *Croton tiglium* seeds (Saputera et. al. 2006). Santha et al. in their study revealed the presence of α-santalol as chemopreventive agent appears to be very promising in skin cancer control (Sreevidya and Chandradhar 2015). The present study also confirms the anti-cancerous activity of *S. album* through the presence of 9 Octadecenoic Acid and Sentlyl acetate in the methanolic extract.

The results pertaining to GC-MS analysis of the methanolic extract of *S. album* lead to the identification of various compounds which have manifold therapeutic benefits. The identified metabolites were known to possess various bioactivities such as antifungal, anti-inflammatory, skin disorders, antibacterial etc. The extract showed the presence of a variety of fatty acids, anti-inflammatory compounds such as hexadecanoic acid, fragrance and flavoring agents such as tetradecanoic acid are found in the plant extract (Burdock and Carabin 2007).

This study explored the therapeutic significance of the sandalwood which has great effect on bacteria causing the metabolite regulation and can be proposed as an herb of psychopharmacological importance. Various chemical compounds which are identified in methanolic extract of sandalwood, *E. coli* lysate and *E. coli* grown in methanolic extract of sandalwood help in the determination of the possible synergistic benefits of the plant and their potential as antimicrobial agent. This also opens up the ways to investigate it further by isolating the single bioactive compounds leading to the further investigation of specific biological and pharmacological potentials (Sermakkani and Thangapandian 2012).

## 5. Conclusions

This study evaluated the antibacterial potential of SwME and the profiling of bioactive compounds generated by GC-MS analysis in SwME extract and in the lysate of *E. coli.* The results concluded that SwME was strong antimicrobial agent showing fair antibacterial activity against all the bacterial strains. GC-MS chromatogram also showed various compounds in SwME with maximum peak intensity which could contribute to the medicinal value of the sandalwood. This study was performed to analyze the effect of sandalwood methanolic extract on the bacterial metabolites when grown in presence of extract. The result showed the presence of compound in the extract such as hexadecanoic acid and 9 Octadecenoic acids which exerts the antimicrobial activity. Antibacterial activity of methanolic extract was found to be impressive against *E. coli* proved that sandalwood has great potential for development of therapeutic agents against microbial ailments and it can also be used as alternative or antibiotic. However, isolation and purification of single bioactive compound and their biological activity will be certainly giving productive outcomes and will open a new area of investigation of individual components and their pharmacological potency. From these results, it could be inferred that *“Santlum album”* contains diverse bio-active compounds. Evaluation of pharmacological activity is under progress. Along these lines, it is suggested as a plant of phyto-pharmaceutical importance.

## Acknowledgements

The authors gratefully acknowledge Advanced Instrumentation Research Facility, JNU, New Delhi for providing the GC-MS facility. We also thank to Department of Biotechnology, Jamia Millia Islamia, and New Delhi for providing the infrastructure to carry out research work.

